# Epithelium intrinsic vitamin A signaling co-ordinates pathogen clearance in the gut via IL-18

**DOI:** 10.1101/667998

**Authors:** Namrata Iyer, Mayara Grizotte-Lake, Sarah R. Gordon, Ana C. S. Palmer, Crystle Calvin, Shipra Vaishnava

## Abstract

Intestinal epithelial cells (IECs) are at the forefront of host-pathogen interactions, coordinating a cascade of immune responses to confer protection against pathogens. Here we show that IEC-intrinsic vitamin A signaling restricts pathogen invasion early in the infection and subsequently activates immune cells to promote pathogen clearance. Mice blocked for retinoic acid receptor (RAR) signaling selectively in IECs (stop^ΔIEC^) showed significantly higher *Salmonella* burden in colonic tissues early in the infection associated with higher luminal and systemic loads of the pathogen at later stages. Higher susceptibility of stop^ΔIEC^ mice correlated with attenuated mucosal interferon gamma (IFNγ) production by underlying immune cells. We found that, at homeostasis, the intestinal epithelium of stop^ΔIEC^ mice produced significantly lower amounts of interleukin 18 (IL-18), a potent inducer of IFNγ. Regulation of IL-18 by vitamin A was also observed in a dietary model of vitamin A supplementation. IL-18 reconstitution in stop^ΔIEC^ mice restored resistance to *Salmonella* by promoting epithelial cell shedding to eliminate infected cells and limit pathogen invasion early in infection. Further, IL-18 augmented IFNγ production by underlying immune cells to restrict pathogen burden and systemic spread. Our work uncovers a critical role for vitamin A in coordinating a biphasic immune response to *Salmonella* infection by regulating IL-18 production by IECs.

**Author Summary:** Epithelial cells line the intestinal lumen, forming a barrier between the body and dietary and microbial contents in the lumen. Apart from absorbing nutrients from diet, these epithelial cells help mediate a stable, symbiotic relationship between microbes in the gut and the immune cells. During infection, they help co-ordinate the immune response to counter the infection. How dietary micronutrients, such as vitamin A, inform epithelial cell function during infection is poorly understood. Using a model where epithelial cells in the gut cannot respond to vitamin A signals, we find that epithelial vitamin A signaling promotes resistance to *Salmonella* infection. We show that, vitamin A increases the production of a key cytokine, interleukin 18, by epithelial cells. IL-18 promotes shedding of infected epithelial cells to reduce the pathogen invasion while also inducing the production of interferon gamma by immune cells to mediate pathogen clearance. Thus, epithelial cells dynamically respond to dietary vitamin A to regulate interleukin 18 production and potentiate resistance to infection.

## Introduction

Resistance to an invasive pathogen involves coordination between the early and late phase of the immune response so as to achieve pathogen clearance without excess collateral damage to the host. The intestinal epithelium is at the forefront of host-microbial interactions and is critical for orchestrating these immune responses during infection. Chemokines secreted by the epithelium are responsible for immune cell recruitment and activation [1]. T cells, NK cells as well as neutrophils are recruited to the colon during infection and secrete pro-inflammatory cytokines such as interferon gamma (IFNγ) to promote bacterial clearance and halt systemic spread of the infection [2, 3]. Moreover, the epithelium itself undergoes cell shedding in the early stages of infection as an innate defense mechanism to clear intracellular pathogens [4, 5]. Mechanisms that orchestrate such diverse functions of intestinal epithelial cells (IECs) during an infection remain poorly studied.

Vitamin A is an important dietary nutrient absorbed in the form of carotenoids and retinyl esters by intestinal epithelial cells and metabolized into its active form retinoic acid (RA). The heterodimeric nuclear receptor complex, retinoic acid receptor (RAR) and retinoid X receptor (RXR), is activated by RA binding to induce target gene expression [6]. Retinoic acid signaling is a potent regulator of gut immune make-up, affecting both the recruitment as well as activity of dendritic cells [7], T cells [8], B cells [9] and innate lymphoid cells (ILCs) [10] in the gut. The retinoic acid synthesized by epithelial cells, while available to underlying immune cells, is also capable of initiating a signaling response within IECs themselves. Retinoic acid signaling in intestinal epithelial cells (IECs) regulates epithelial lineage specification and promotes small intestinal T helper 17 (Th17) responses [11, 12]. Dietary vitamin A deficiency markedly increases susceptibility to enteric pathogens [13], however, relatively little is known about the contribution of IEC-intrinsic retinoic acid signaling in the context of infection.

In this study we use a mouse model expressing IEC-specific dominant negative retinoic acid receptor (stop^ΔIEC^) to investigate the role of retinoic acid signaling during infection. stop^ΔIEC^ mice are more susceptible to luminal and systemic colonization by *Salmonella*. This is associated with abrogated shedding of infected epithelial cells as well as a blunted interferon gamma (IFNγ) response. We find that expression of interleukin-18, a known inducer of IFNγ, is dependent on retinoic acid signaling in intestinal epithelial cells. RAR signaling-dependent IL-18 promotes epithelial cell shedding as well as mucosal IFNγ production to orchestrate resistance to infection. Our results thus reveal a novel regulatory axis in the gut, wherein epithelial-intrinsic signaling in response to vitamin A, sequentially triggers IL-18 dependent mechanisms to first limit tissue invasion and then trigger an IFNγ response to promote pathogen clearance.

## Results

### Epithelial-intrinsic RAR signaling is protective against *Salmonella* colonization

Vitamin A deficiency results in increased susceptibility to infection by *Salmonella* and other enteric pathogens [13, 14]. Vitamin A deficiency causes immune dysregulation in the gut, including improper lymphoid recruitment, maturation and functional potential [15, 16]. These sweeping changes obscure the finer details of how vitamin A regulates infection outcome. Dietary vitamin A is sequentially absorbed, metabolized and distributed by intestinal epithelial cells [17]. Retinoic acid synthesized by intestinal epithelial cells regulates immune cell recruitment and cytokine production [18, 19]. We therefore hypothesized that vitamin A signaling in IECs regulates the mucosal immune response to mediate infection susceptibility. To address this question, we chose a mouse model wherein vitamin A signaling was specifically abrogated in intestinal epithelial cells while metabolism of vitamin A was unchanged. This was done through the Villin-Cre dependent overexpression of a dominant negative form of retinoic acid receptor alpha [20, 21], found to be the most dominant isoform expressed in intestinal epithelial cells (Fig 1A). Mice overexpressing this dominant negative RAR (stop^ΔIEC^) were defective in the expression of the RA-responsive gene *isx* compared to their wild type littermates (stop^flox^) (Fig 1B and 1C) [22]. At homeostasis, stop^ΔIEC^ mice displayed an increase in goblet cell differentiation as well as overall thickness of the mucus barrier (S1A and S1B Fig), corroborating the results of a study using RARa^ΔIEC^ mice [11]. However, stop^ΔIEC^ mice showed no significant differences in epithelial permeability or turnover (S1C and S1D Fig).

**Fig 1.**
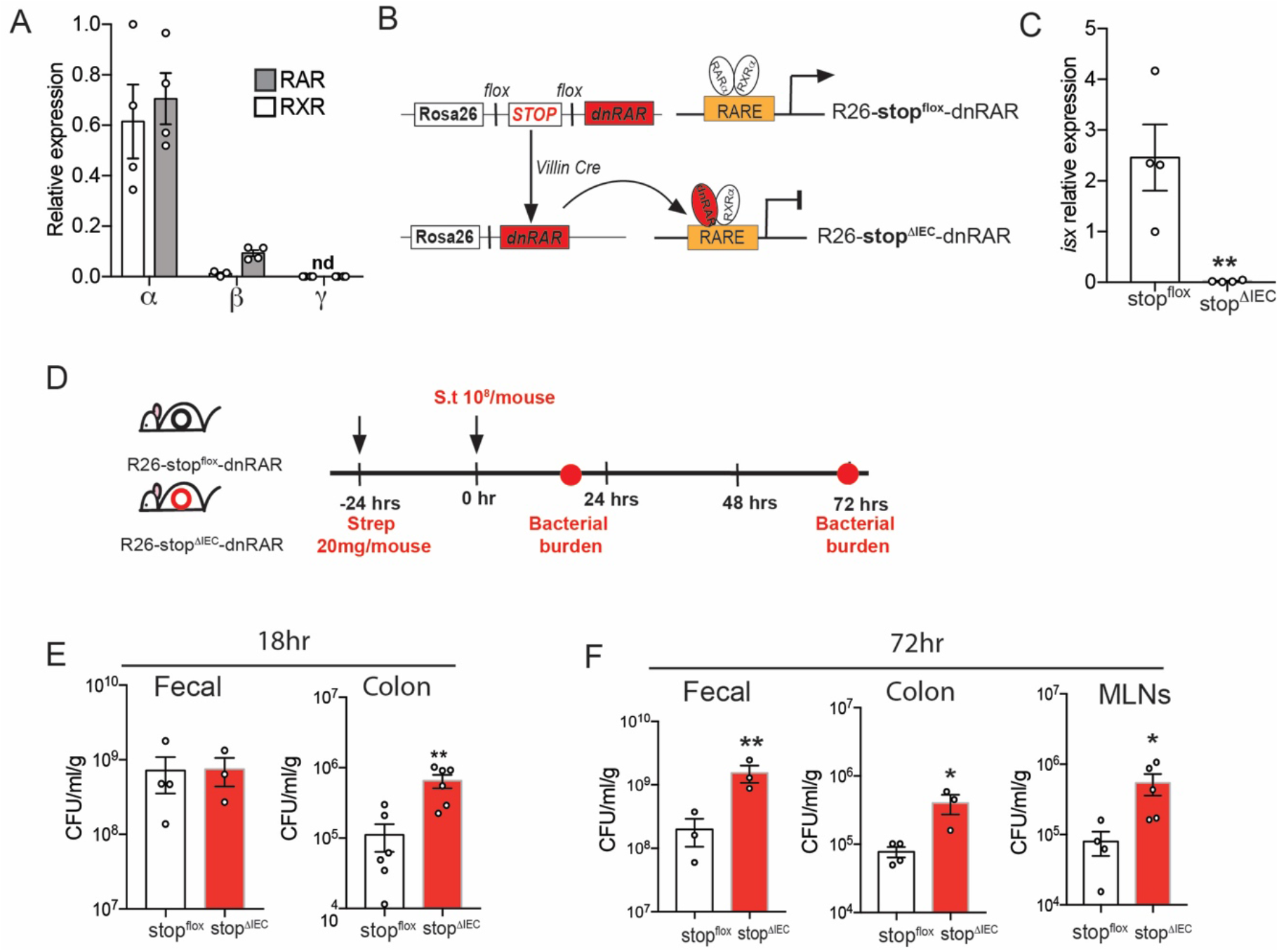
Epithelial-intrinsic RAR signaling is protective against *Salmonella* colonization. **(A)** Relative expression of RAR and RXR isoforms (α, β and γ) in laser capture microdissected epithelial cells from homeostatic C57BL/6 mice ileal tissues **(B)** Schematic representation of the creation of stop^flox^ (wild type) and stop^ΔIEC^ (RAR signaling knockdown) mice using Villin-Cre dependent expression of the dnRAR cassette. **(C)** Relative expression of vitamin A responsive gene, *isx*, in stop^flox^ and stop^ΔIEC^ ileum tissues. **(D)** Schematic representation of *Salmonella* infection timeline with assessment of pathogen loads at 18 hours and 72 hours post infection (hpi). **(E)** Bacterial burden in fecal and proximal colon tissues at 18 hpi in stop^flox^ and stop^ΔIEC^ mice. **(F)** Bacterial burden in fecal, distal colon and mesenteric lymph node samples at 72 hpi in stop^flox^ and stop^ΔIEC^ mice. Representative data from 3 independent experiments. n=3-4 mice per group. Student’s t test was used for statistical analysis. *P<0.05; **P<0.01.

The role of epithelial-intrinsic vitamin A signaling during infection was assessed using a gastroenteritis infection model of non-typhoidal *Salmonella* Typhimurium (Fig 1D) [23]. Bacterial burden in the feces, colon and mesenteric lymph nodes was determined at early (18 hours post infection; hpi) and late (72 hpi) time points to assess if vitamin A signaling modulates the kinetics of the pathogen colonization. At early time points, stop^ΔIEC^ mice and stop^flox^ littermate controls had similar luminal burdens of the pathogen. However, stop^ΔIEC^ mice showed significantly higher loads of the bacterium within colon tissues (Fig 1E). This advantage in initial tissue invasion bolstered pathogen burdens at later time points, with higher loads found in the feces, colon as well as mesenteric lymph nodes of stop^ΔIEC^ mice (Fig 1F). These results show that loss of RAR signaling in the intestinal epithelium phenocopies dietary vitamin A deficiency in the context of enteric infection. Further, the changes in infection kinetics suggest that epithelial-intrinsic vitamin A signaling co-ordinates early and late immune mechanisms to promote resistance to infection.

### Epithelial-intrinsic RAR signaling promotes mucosal IFNγ response during infection

Vitamin A metabolism and signaling in the epithelium is a key determinant of the intestinal immune make up. In the gut, retinoic acid influences the balance between Th1 and Th17 cells, both of which are important for controlling infection [8, 24, 25]. IEC-intrinsic vitamin A metabolism promotes mucosal IL-22 production, which in turn induces dysbiosis and aids pathogen colonization [18]. We therefore analyzed the colonic lamina propria populations in our mouse model to check if the increased susceptibility of stop^ΔIEC^ mice is due to a dysregulated immune response. While stop^ΔIEC^ and stop^flox^ mice showed similar colonic immune make-up at homeostasis (S2A-S2D Fig), on day 3 of *Salmonella* infection, stop^ΔIEC^ mice displayed a defect in interferon gamma production (Fig 2A). CD4 and CD8 T cells were recruited to the colon to similar extents in stop^ΔIEC^ and stop^flox^ mice, however activation of these cells to produce IFNγ was defective in stop^ΔIEC^ mice (Fig 2B-2D). No significant differences in mucosal IL-17 and IL-22 production were observed between stop^ΔIEC^ and stop^flox^ mice during infection (S3A and S3B Fig).

**Fig 2.**
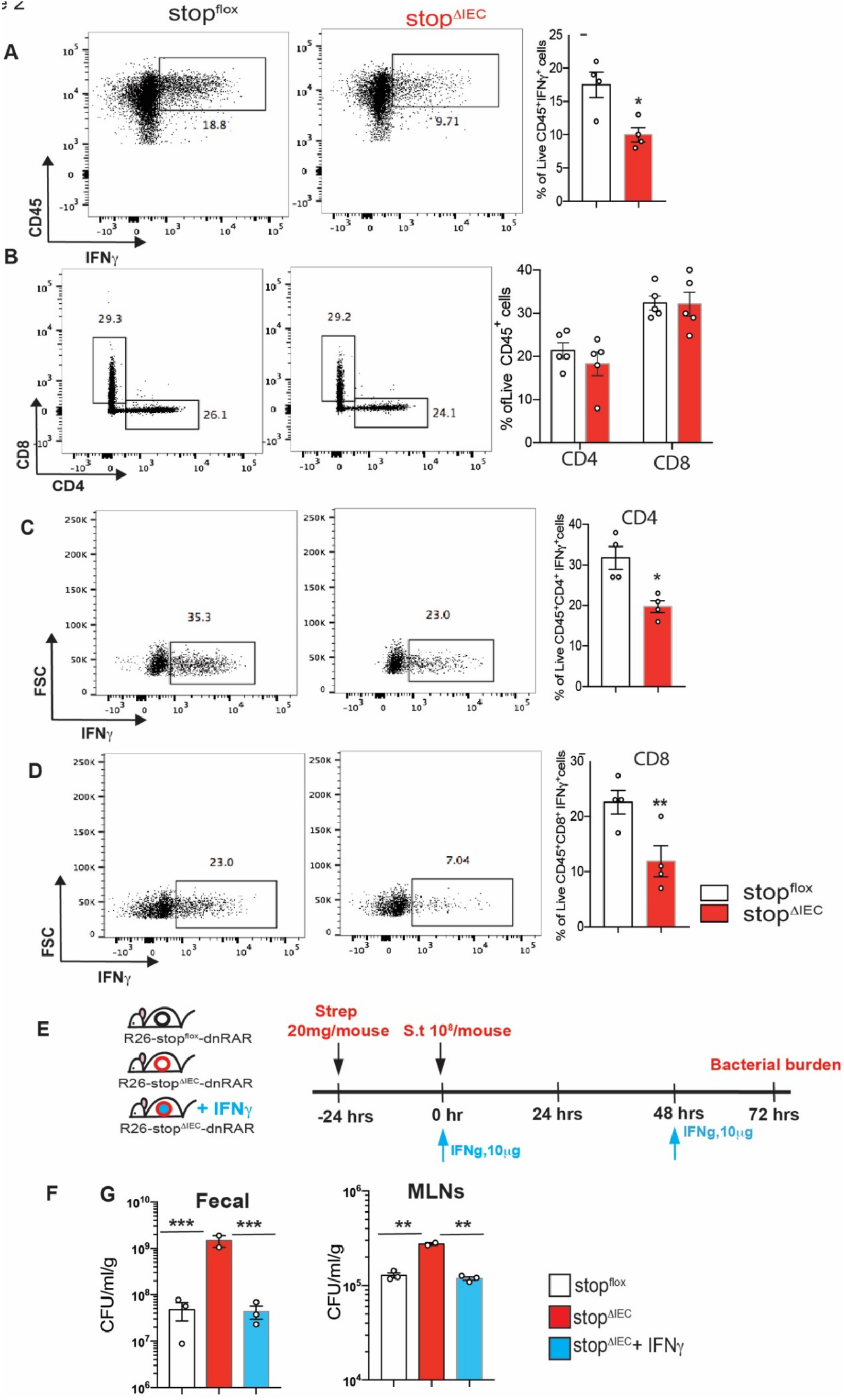
Epithelial-intrinsic RAR signaling promotes mucosal IFNγ response during infection. **(A-D)** Flow cytometry analysis of colonic lamina propria lymphocytes from stop^flox^ and stop^ΔIEC^ mice 72 hpi with *Salmonella*. Representative density plots and quantitative analysis of relative frequencies of **(A)** total live CD45+ IFNγ+ cells **(B)** total CD4+ and CD8+ T cells **(C)** CD4+ IFNγ+ cells and **(D)** CD8+ IFNγ+ cells in stop^flox^ and stop^ΔIEC^ mice. Representative data from 2 independent experiments. n=3-4 mice per group. Student’s t test was used for statistical analysis. *P<0.05; **P<0.01. **(E)** Schematic representation of IFNγ feedback in stop^ΔIEC^ mice during *Salmonella* infection. Mice were intraperitoneally injected with 10 μg of IFNγ at 0 hpi and 48 hpi. **(F)** Bacterial burden in fecal and mesenteric lymph nodes at 72 hrs post *Salmonella* infection in stop^flox^, stop^ΔIEC^ and stop^ΔIEC^ + IFNγ mice. One-way ANOVA was used for statistical analysis. **P<0.01, ***P<0.005.

Previous studies, both *in vitro* and *in vivo*, have shown that interferon gamma can be regulated by vitamin A, with retinoic acid promoting IFNγ production in intestinal T cells [26, 27]. Interferon gamma plays an important role in resistance to *Salmonella* infection. IFNγ knockout mice are more susceptible to *Salmonella*, both at intestinal and systemic level [28]. Mice with distinct microbiome compositions were found to be differentially susceptible to *Salmonella*; a function of the IFNγ induction by these communities [2]. Mechanistically, IFNγ promotes phagocytosis, oxidative/nitrosative response and cellular autophagy to clear the pathogen [29, 30]. To assess if the defect in IFNγ production was responsible for the increased susceptibility in stop^ΔIEC^ mice, we performed IFNγ feedback experiments (Fig 2E). Reconstitution of IFNγ in stop^ΔIEC^ mice led to a rescue of susceptibility, with bacterial burdens returning to wild type levels (Fig 2F). These results suggest, that epithelial-intrinsic RAR signaling primes mucosal IFNγ production to restrict *Salmonella* infection.

### Intestinal epithelium-intrinsic RAR signaling regulates interleukin-18

Interleukin-18 was first discovered as an interferon gamma inducing molecule [31, 32]. Upon transcription and translation, it is present within the cell as a precursor form (pro-IL18) which is proteolytically processed by caspases into its mature, secreted form [33]. This IL-1 family cytokine, unlike its close cousin IL 1β, is constitutively expressed by a wide variety of cell types in the body. In the gut, intestinal epithelial cells form the main source of IL-18 [34, 35]. IL-18 is microbially regulated. It accumulates in the intestine after microbial colonization and is responsive to microbial metabolites such as butyrate and taurine [36, 37]. Further, IL-18 is known to be up-regulated at the early stages of *Salmonella* infection [3]. A study with human neuroblastoma cells identified induction of IL-18 by all trans retinoic acid *in vitro* [38]. Further, serum IL-18 levels have been shown to increase during vitamin A supplementation in obese mice; an effect dependent on active vitamin A metabolism [39]. However, regulation of homeostatic IL-18 levels in the gut by vitamin A has not been previously reported. We hypothesized that interleukin-18 might be the mechanistic link between IEC-intrinsic RAR signaling and mucosal IFNγ response.

We assessed the levels of IL-18 at homeostasis between stop^ΔIEC^ and stop^flox^ mice. Colon whole tissue (Fig 3A) as well as colonocytes (Fig 3B) of stop^ΔIEC^ mice showed reduction in the protein levels of the precursor form of IL-18. These results suggest that RAR signaling in intestinal epithelial cells promotes *il18* gene expression in the gut. In order to confirm that this phenomenon is not restricted to our stop^ΔIEC^ mouse model, we confirmed the same in a dietary model where wild type mice were fed a diet spiked with retinyl acetate for 3 days. Compared to mice receiving vehicle control, mice fed excess retinyl acetate had increased levels of IL-18 in colonocytes (Fig 3C). These results suggest that dietary vitamin A can dynamically modulate the levels of IL-18 in the colon.

**Fig 3.**
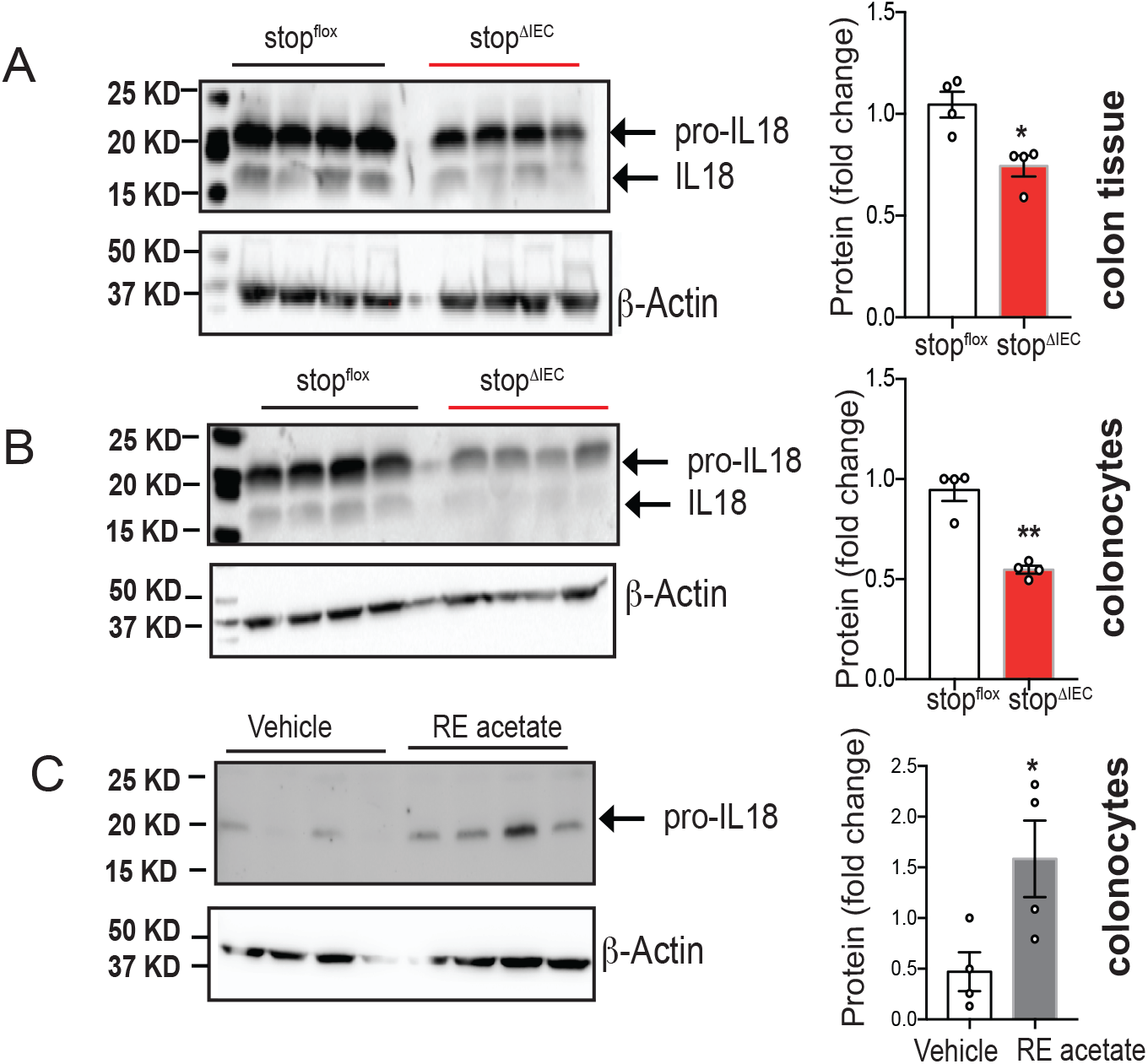
Intestinal epithelium-intrinsic RAR signaling regulates interleukin-18. Representative western blots and quantitative image analysis results comparing homeostatic levels of IL-18 in colon whole tissue **(A)** and colonocyte **(B)** lysates from stop^flox^ and stop^ΔIEC^ mice. Representative data from 2 independent experiments. **(C)** Representative western blot and quantitative image analysis comparing IL-18 levels in colonocytes from stop^flox^ mice fed regular mouse chow spiked with vehicle (corn oil) or retinyl acetate (500 IU/g) for 3 days. Representative data from 2 independent experiments. All quantitation analysis done for pro-IL18 levels in all experiments. ImageJ was used for densitometric analysis of image. β actin levels were used for normalization. Student’s t test was used for statistical analysis. *P<0.05; **P<0.01.

### RAR signaling-dependent IL-18 orchestrates early resistance to *Salmonella* invasion

Our results in stop^ΔIEC^ mice demonstrated a direct correlation between intestinal IL-18 levels and resistance to *Salmonella* infection. In order to establish a causal relationship between IL-18 and infection outcome, we reconstituted IL-18 levels in stop^ΔIEC^ mice and compared their susceptibility to stop^flox^ and stop^ΔIEC^ mice at early time points (18 hpi) of *Salmonella* infection. We found that while IL-18 left luminal colonization unaffected, IL-18 feedback in stop^ΔIEC^ mice completely rescued tissue burdens of the pathogen to levels lower than wild type (Fig 4A-4C). This was further confirmed using confocal microscopy where staining for *Salmonella* revealed significantly higher levels of bacteria within stop^ΔIEC^ colonic tissues compared to stop^flox^ and IL-18 feedback mice (Fig 4D-4F; S1-S3 Movies). Feedback with IFNγ, however, failed to rescue early tissue invasion in stop^ΔIEC^ mice (Fig 4A-4C). This suggests that epithelial RAR signaling promotes early resistance to bacterial invasion in an IL-18 dependent, but IFNγ independent, manner.

**Fig 4.**
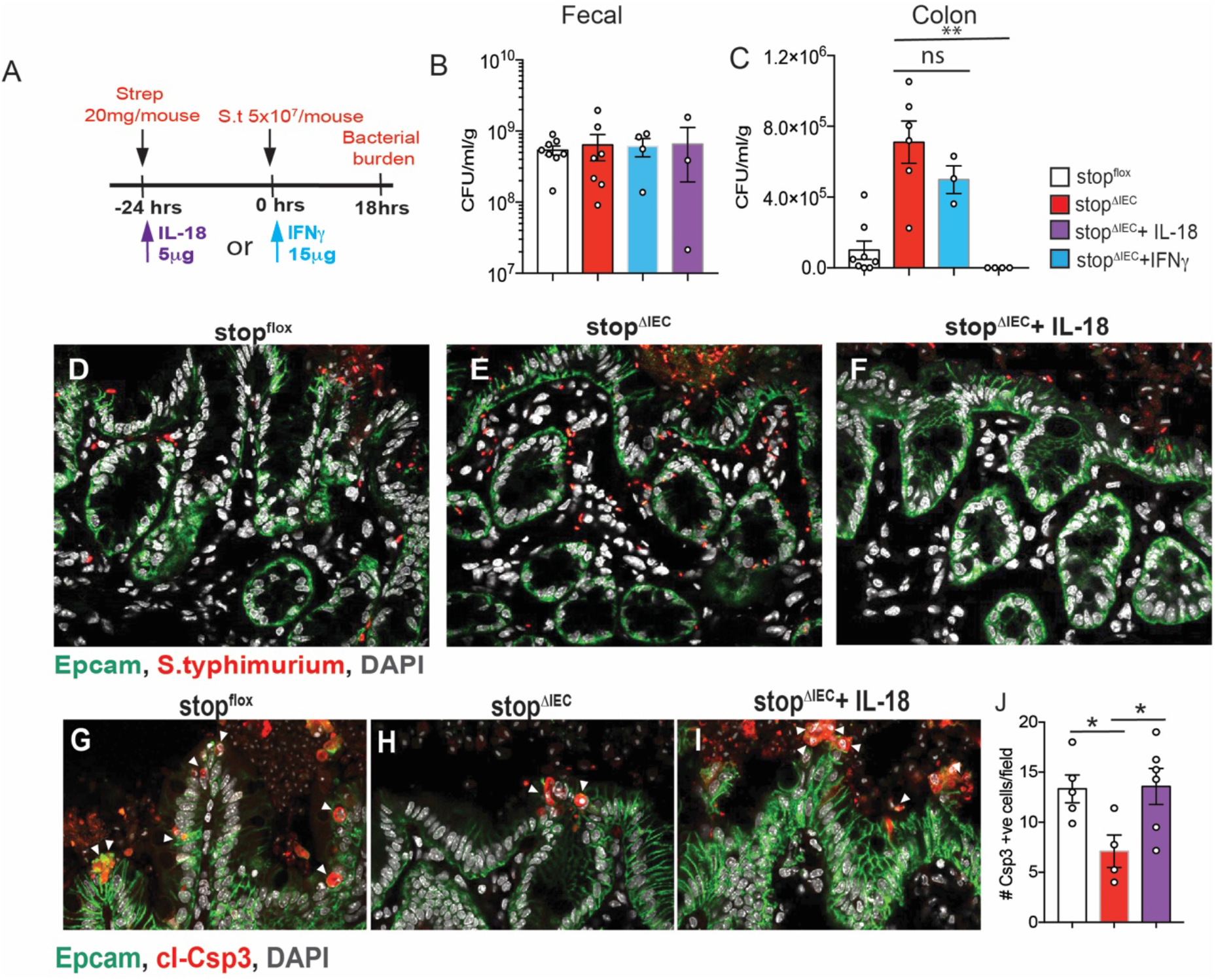
RAR signaling-dependent IL-18 orchestrates early resistance to *Salmonella* invasion. **(A)** Schematic representation of IL-18 and IFNγ feedback regimen for 18 hour time point in *Salmonella* infection. **(B)** Fecal and **(C)** Colon bacterial loads at 18hpi in stop^flox^, stop^ΔIEC^, stop^ΔIEC^ + IL-18 and stop^ΔIEC^ + IFNγ mice. Combined data from 2 independent experiments **(D, E and F)** Representative confocal microscopy images depicting loads of *Salmonella* (red) in colon tissue 18 hrs post infection, counterstained with EpCAM (epithelial cells; green) and DAPI (nuclei; grey). **(G, H and I)** Representative images and **(J)** quantitative analysis of epithelial cell death (cleaved caspase-3 positive cells; red) in cecal tissues of stop^flox^, stop^ΔIEC^ and stop^ΔIEC^ + IL-18 mice at 18 hours post infection. Samples counterstained with Epcam (epithelial cells; green) and DAPI (nuclei; grey). Combined data from 2 independent experiments. Quantitative comparison was made by counting total Csp3+ve cells per image. Data is an average of 6-10 images per mouse with 3-4 mice per group per experiment. Oneway ANOVA was used for statistical analysis. *P<0.05; **P<0.01, ***P<0.005.

The epithelium is the first point of contact with adherent and invasive pathogens. Early in the infection, *Salmonella* uses epithelial invasion as a strategy to induce tissue inflammation, which endows it a selective advantage over gut commensals [40, 41]. Restricting this invasion is important to limit luminal expansion and systemic spread of the pathogen. Recently, epithelial cell death has been identified as a strategy to eliminate infected cells in the gut and limit tissue colonization. While cell shedding routinely occurs as a part of epithelial turnover, the rate of shedding increases dramatically during pathological insult to the epithelium [42, 43]. In the context of infection, inflammasome-mediated cell death pathways have been implicated in this epithelial-intrinsic response [4, 44]. The activation of these cell death pathways is associated with IL-18 cleavage and secretion [42]. We hypothesized that RAR signaling in IECs modulates cell death response during *Salmonella* infection via IL-18. Cell death has been reported to occur as an early response to the infection, especially in the cecum which is more permissive to bacterial invasion [4]. We, therefore, assessed the extent of cecal epithelial cell shedding in stop^ΔIEC^ mice at 18 hpi using cleaved caspase-3 staining as a marker for dying epithelial cells. We observed significantly higher numbers of dying cells in stop^flox^ mice compared to stop^ΔIEC^ mice. This defect was rescued completely upon IL-18 reconstitution (Fig 4G-4J). This suggests that RAR signaling mediated IL-18 production promotes an epithelial cell shedding response which is associated with restricted tissue invasion by *Salmonella*.

### Intestinal epithelium-intrinsic RAR signaling regulates pathogen colonization via interleukin-18

Our results with IFNγ feedback showed that at day 3 of infection, IFNγ is successfully able to rescue pathogen colonization in stop^ΔIEC^ mice (Fig 2F). IL-18, but not IFNγ, feedback was required to rescue early tissue invasion susceptibility in stop^ΔIEC^ mice (Fig 4C). Therefore, we hypothesized that IL-18 levels in the gut promoted early innate defenses to infection, while priming mucosal IFNγ to mediate pathogen clearance in the later stages. To evaluate this, we analyzed the effect of IL-18 reconstitution in stop^ΔIEC^ mice on day 3 of infection (Fig 5A). Feedback of IL-18 was sufficient to bring down bacterial loads in fecal, MLN as well as colon samples of stop^ΔIEC^ mice back to wild type levels (Fig 5B and 5C). Assessment of the colonic lamina propria revealed that this feedback effectively rescued IFNγ production by mucosal T cells (Fig 5D and 5E). These results suggest that epithelial RAR signaling modulates the mucosal IFNγ response via IL-18 to mediate resistance to pathogen. These results align with a study where epithelial IL-18 expression, regulated by histone deacetylase 3 activity, primes IFNγ response in intraepithelial lymphocytes to restrict colonization by *Citrobacter* [45]. IL-18 induction by protozoan colonization in the gut exerts a similar protective effects against *Salmonella* by promoting mucosal Th1 and Th17 response [46]. Our results corroborate the importance of IL-18 as a determinant of infection susceptibility and reveal a previously unappreciated pathway for IL-18 regulation via vitamin A signaling and dietary vitamin A.

**Fig 5.**
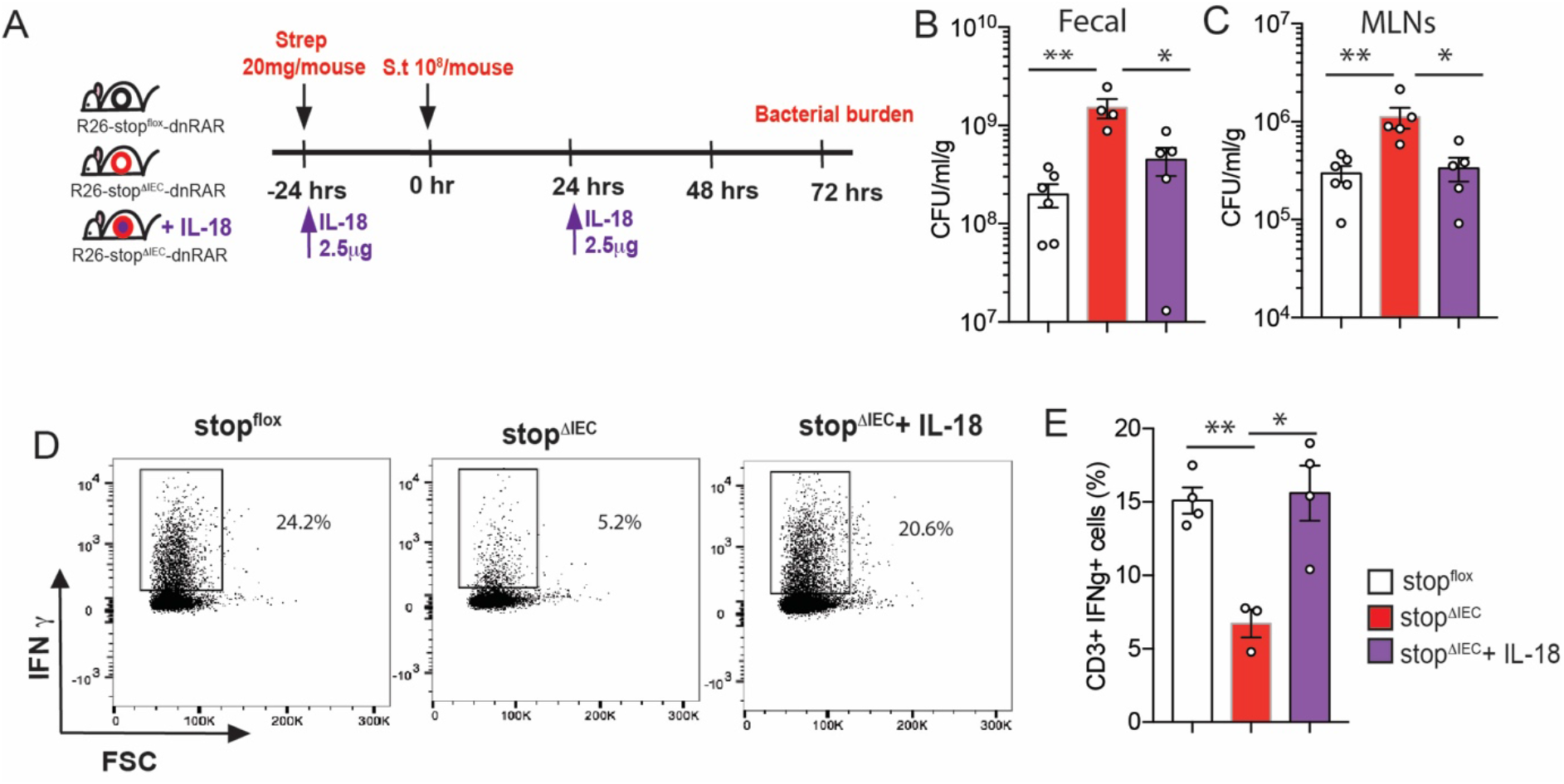
Intestinal epithelium-intrinsic RAR signaling regulates pathogen colonization via interleukin-18. **(A)** Schematic representation of IL-18 feedback regimen in stop^ΔIEC^ mice during *Salmonella* infection at 72 hpi. Bacterial burden in fecal **(B)**, mesenteric lymph nodes **(C)** and colon **(D)** at 72 hpi in stop^flox^, stop^ΔIEC^ and stop^ΔIEC^ + IL-18 mice. Combined data from 2 independent experiments. **(E)** Flow cytometry analysis of colonic lamina propria lymphocytes from stop^flox^, stop^ΔIEC^ and stop^ΔIEC^ + IL-18 mice with density plots and quantitative analysis of relative frequencies of CD3+IFNγ+ cells. Representative data from 2 independent experiments. n=3-4 mice per group. One-way ANOVA was used for statistical analysis. *P<0.05; **P<0.01, ***P<0.005.

## Discussion

Vitamin A is a potent dietary micronutrient and vitamin A deficiency causes susceptibility to a spectrum of infectious diseases, especially in developing countries. A large body of work has contributed to our understanding of the immunomodulatory potential of vitamin A and its metabolite retinoic acid. However, pleiotropic cellular effects, complex metabolic and distribution pathways as well as source- and concentration-dependent functional effects have made dietary vitamin A models of infection difficult to interpret [47]. In this study, we employ a tissue-specific signaling abrogation model to elucidate the specific role of the vitamin A signaling pathway in the intestinal epithelium during infection. A key feature of this model is the absence of interference with the vitamin A metabolic machinery, avoiding any differences in gut immune trafficking mediated by epithelium-derived RA. Specifically, this study extends our understanding of homeostatic functions of vitamin A signaling in the intestine and reveals a previously unappreciated regulatory role for this pathway during *Salmonella* infection in the gut. In addition to describing the dynamic regulation of homeostatic IL-18 levels in the gut by vitamin A, we elucidate the functional role of the vitamin A-IL18 axis in restricting early tissue invasion by *Salmonella* and priming mucosal IFNγ production to mediate pathogen clearance (Fig 6).

**Fig 6.**
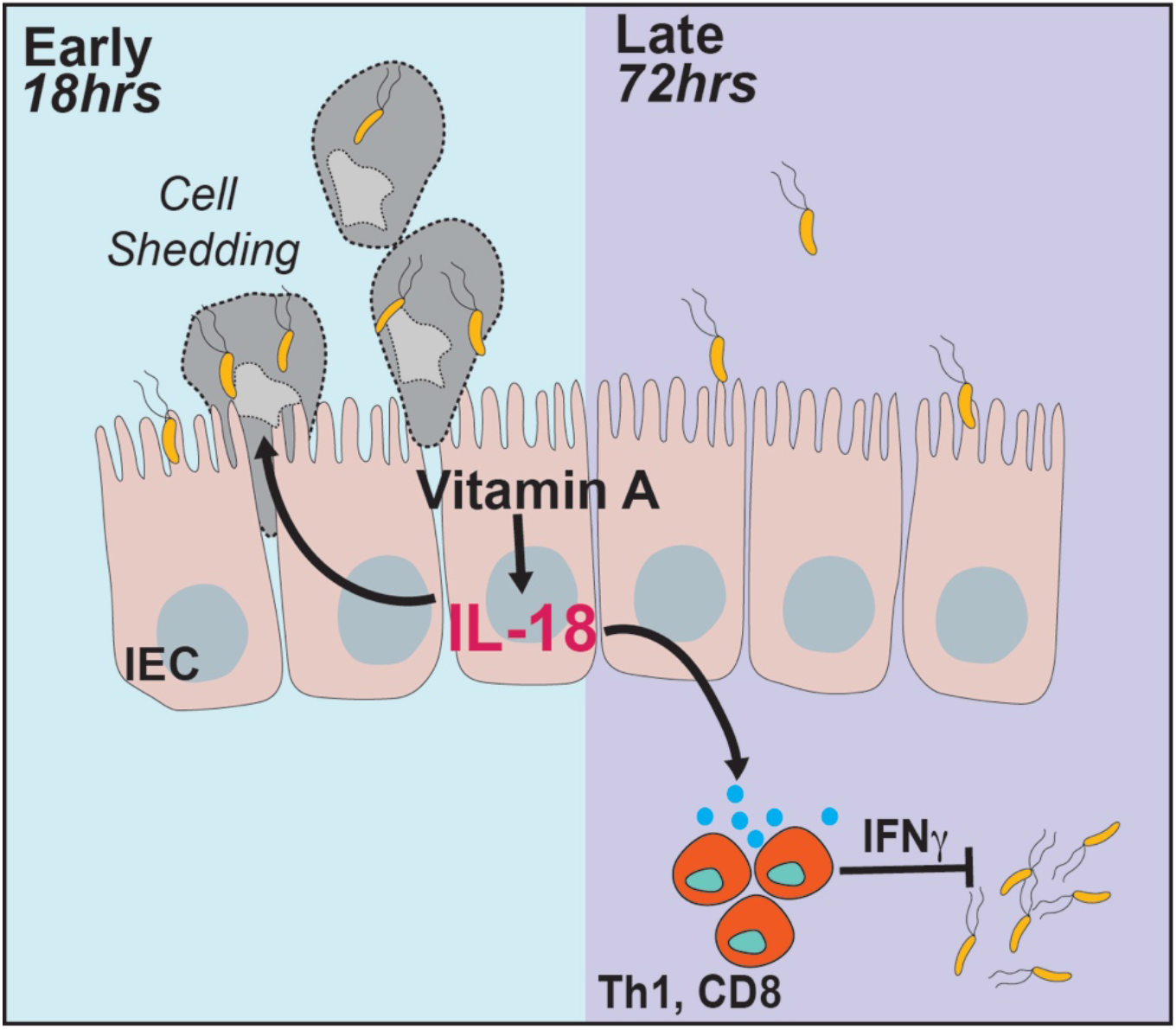
Model depicting role of epithelial-intrinsic signaling during *Salmonella* infection. Dietary vitamin A activates retinoic acid signaling within colonic epithelial cells to induce production of IL-18 at homeostasis. During early stages of *Salmonella* infection (18 hours), IL-18 promotes epithelial cell shedding to eliminate infected cells and restrict pathogen invasion. IL-18 also primes IFNγ production by mucosal T cells which promote pathogen clearance during later stages of the infection (72 hours).

Cytokines secreted by epithelial cells are important for immune co-ordination in the gut. Vitamin A is known to induce the signaling of important cytokines such as IL-22 in the gut [26]. A previous study from our group has shown that epithelial-intrinsic retinoic acid synthesis promotes IL-22 expression in underlying immune cells which in turn promotes *Salmonella* induced dysbiosis and gut colonization [18]. In contrast, the current study shows that epithelial-intrinsic RA signaling regulates IL-18 expression which protects against pathogen colonization and systemic spread. This highlights that retinoic acid synthesized by epithelial cells has distinct autocrine and paracrine functions, each having differential effects on infection outcome.

Interleukin-18 is set apart from other members of the IL1 family by being constitutively expressed by wide range of cell types throughout the body [48]. While both IL1β and IL-18 have pro-inflammatory functions, the classic fever response associated with IL1β is absent with IL-18. TLR ligands (LPS, poly (I:C), pam3CSK4) and type I interferons induce expression of IL-18, which is then proteolytically processed via the inflammasome pathway [49]. While induction of IL-18 during inflammatory conditions is well documented, relatively little is known about homeostatic regulators of IL-18. Studies have shown that late in gestation, inactive IL-18 starts accumulating in the intestine and active IL-18 is detectable postnatally [50]. Microbial colonization of the gut induces an upregulation in IL-18 expression [37] and the microbial metabolite butyrate is implicated in transcriptional regulation of IL-18 [36]. This study is the first to demonstrate that homeostatic IL-18 levels in the gut are regulated by IEC-intrinsic vitamin A signaling. Further, we show that vitamin A supplementation in the diet is also capable of inducing IL-18 in the gut, suggesting a dynamic regulation of epithelial IL-18 levels by dietary vitamin A.

Although our results clearly demonstrate that IL-18 levels in the gut are regulated by vitamin A, the exact mechanism of this regulation remains unresolved. Studies mining transcriptional targets of RAR signaling by *in silico* and ChIP-seq techniques have not identified IL-18 as a candidate, suggesting that retinoic acid receptor does not directly bind the *il18* promoter to induce its expression [51, 52]. IL-18 expression is regulated by a host of transcription factors such as NF-κB, PU.1, Stat1, AP-1 and Bcl6 [49, 53, 54]. Both Stat1 and Bcl6 are reported to be regulated by retinoic acid [55–57]. Further, a post-transcriptional regulators of IL-18, Bruton’s tyrosine kinase, also has a predicted RAR binding site [58, 59]. We hypothesize that vitamin A regulation of IL-18 occurs by an indirect mechanism involving an interplay between one or more direct targets of RAR signaling. Uncovering the mechanistic link between vitamin A and IL-18 will be the focus of future investigations.

IL-18 and IFNγ are implicated in a host of gastrointestinal and systemic infections caused *Toxoplasmas gondii* [60], *C. rodentium* [45], *Listeria monocytogenes* [61, 62] among others. These pathogens represent a diversity of adhesion/invasion strategies as well as potential for systemic spread and toxicity. It remains to be seen how epithelial-intrinsic vitamin A signaling and IL-18 modulate infection outcome in these contexts.

## Materials and Methods

### Mice

All mice were bred in the SPF barrier facility at Brown University. Wild type stop^flox^ mice were a kind gift from Dr. S. Sockanathan (Johns Hopkins University School of Medicine). Villin-Cre mice in a C56BL/6J background were purchased from Jackson Laboratories. stop^flox^ mice were bred to Villin-Cre mice to get stop^flox^ (Cre negative) or stop^ΔIEC^ (Cre positive) mice. All mice used in the study were 6-10 weeks old and were gender-matched across different groups. Littermates or co-housed mice were used to minimize microbiome mediated effects on the study.

### Bacterial strains and maintenance

All infection experiments were carried out using *Salmonella* Typhimurium SL1344 GFP strain (kind gift of Dr. Vanessa Sperandio, UT Southwestern). Strain was routinely maintained on Luria Bertani agar plates containing 100 μg/ml ampicillin.

### Mouse infections

For a gastroenteritis model of *Salmonella* infection protocol outlined by Barthel et al. was followed [23]. Briefly, mice were deprived of food and water for 4 hours and then orally gavaged with 20 mg streptomycin. After 20 hours, mice were again deprived of food and water for 4 hours. Overnight culture of *Salmonella* was subcultured for 4 hours, following which mice were infected with 10^8^ bacteria by oral gavage. Mice were sacrificed 72 hours post infection to harvest organs and assess bacterial burden. For bacterial burden in distal colon, tissues were harvested, fileted and washed twice in phosphate-buffered saline (PBS). Tissues were then incubated in PBS containing 400 μg/ml gentamicin for 30 min at room temperature with shaking. Tissues were then rinsed with PBS, weighed, homogenized and plated to determine bacterial burden.

In order to assess the infection at early time points, protocol outlined by Sellin et al was followed [4]. The streptomycin treatment and *Salmonella* subculture was done as described above and mice were infected with 5×10^7^ bacteria by oral gavage. Assessment of cell death as well as tissue burden were performed in the cecum and proximal colon respectively which are the primary sites infected at this early time point. For cell death assessment, cecal tissues were harvested and fixed in 4% paraformaldehyde/4% sucrose overnight. Fixed tissues were then saturated in PBS containing 20% sucrose, embedded in optimal cutting temperature (OCT) medium, frozen over dry ice and stored at −80°C. Proximal colon tissues were washed and gentamycin-treated for assessment of bacterial burden.

For reconstitution experiments, mouse recombinant IFNγ (Sino Biological; Cat. #50709-MNAH) and mouse recombinant IL-18 (Sino Biological; Cat. # 50073-MNCE) were used. Since IFNγ production defect in stop^ΔIEC^ mice was observed only after infection, reconstitution was performed 0 hr and 48hr post infection with 10 μg recombinant IFNγ intraperitoneally. Since stop^ΔIEC^ mice had homeostatic deficiency in IL-18 production, they were reconstituted with 2.5 μg recombinant IL-18 intraperitoneally 24 hours before and 24 hours after infection. For the early time point experiments, stop^ΔIEC^ mice received a single dose of 5 μg recombinant IL-18 intraperitoneally 24 hours before infection or a single dose of 15 μg recombinant IFNγ intraperitoneally 0 hours post infection.

### Colonic lamina propria lymphocyte isolation and analysis

Colonic lamina propria lymphocytes were isolated as described by Kim et al [63]. Briefly, colons were flushed to remove luminal content, fileted, cut into three pieces and stored in ice-cold PBS. Tissues were washed by vigorous shaking, followed by sequential 10 min digestions at 37°C in HBSS containing 3% FCS, 1mM dithiothreitol and 30mM ethylene diamine tetraacetic acid (EDTA) and HBSS containing 3% FCS and 30mM EDTA. Tissues were vigorously shaken between treatments to dislodge epithelial cells. Lymphocytes were liberated from the lamina propria by digesting tissues in RPMI complete containing 40 μl/ml collagenase (Sigma-Aldrich, stock solution 5 mg/ml) and 5 μl/ml DNase (Sigma-Aldrich, stock solution 0.5 mg/ml) for one hour at 37°C followed by vigorous shaking. Cells were strained through a 70 μm filter and separated on a discontinuous 40%/80% Percoll (GE Healthcare) gradient. Cells at the interface were harvested and processed for flow cytometry analysis.

### Flow cytometry analysis

Cells were stimulated for 4 hours at 37°C in RPMI complete containing 1X cell stimulation cocktail (eBioscience) and protein transport inhibitor cocktail (eBioscience). Stimulated cells were harvested and stained for surface markers and viability for 30 min at room temperature. Samples were stained with viability dye APCef780 (ThermoFisher), CD45 evolve655 (ThermoFisher), CD4 BV785 (BioLegend), CD3 eF450 (ThermoFisher), CD8a BV605 (BioLegend). Following overnight fixation using the Fixation/Permeabilization solution (eBioscience Foxp3 Staining Buffer Set), cells were permeabilized and stained for cytokines IL-17A AF488 (eBioscience), IL-22 PE (eBioscience) and IFNγ PE (eBioscience). Cells were analyzed using the Aria IIIu cytometer and data was analyzed using the FlowJo software.

### Laser capture microdissection and quantitative real time PCR

Intestinal tissues were flushed with PBS and OCT, embedded in OCT, frozen on dry ice and stored at −80°C. Cryosections (10 μm thick) were stained with methyl green and eosin. Intestinal epithelial cells were selectively captured using the Arcturus Laser capture Microdissection system. RNA was isolated from IECs using the RNAqueous-Micro Total RNA kit (Ambion) and converted into cDNA using the iScript cDNA synthesis kit (BioRad). Whole tissue samples were homogenized, RNA was extracted using the PureLink RNA isolation kit (Life Technologies) and converted to cDNA using MMLV reverse transcriptase. Quantitative real time PCR was carried out using SYBR green master mix (Maxima). Expression was normalized using *gapdh* as a housekeeping gene. Details of primers are provided in Table 1.

**Table 1.**
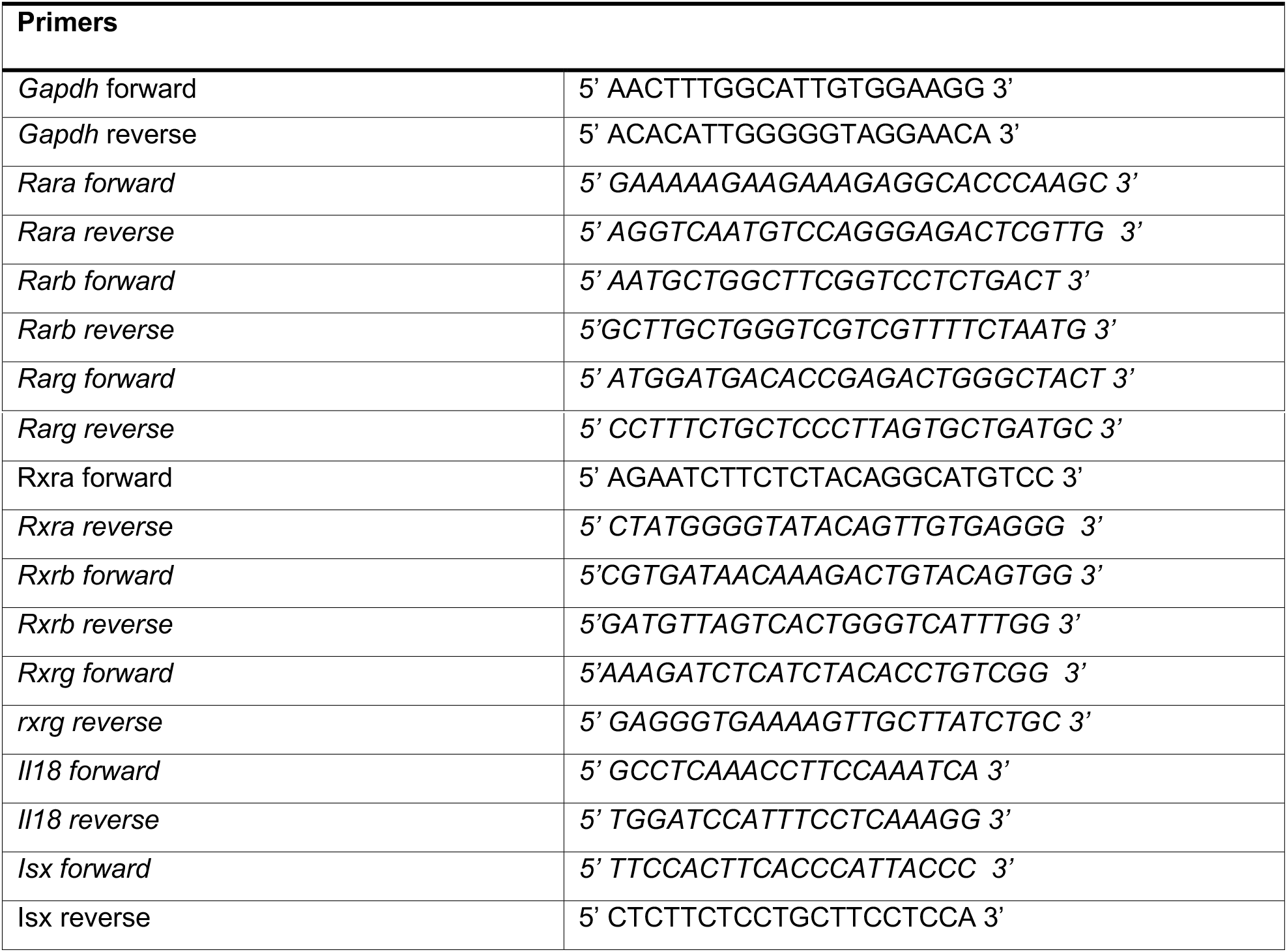
Primers used in this study.

### Barrier function analysis

BrdU incorporation assay was used for assessment of barrier turnover. Mice were injected intraperitoneally with 1 mg of bromodeoxyuridine. Mice were sacrificed 2 hours and 24 hours post injection, distal colon tissues were fixed in methacarn fixative and embedded in paraffin blocks. Tissue sections (7 μm thick) were stained with anti-BrdU antibody (Novus Biologicals; #NBP2-14890) and visualized. For mucus staining, colon tissues samples were fixed in methacarn and embedded in paraffin blocks. Tissue sections (7 μm thick) were stained with alcian blue-periodic acid Schiff’s reagent to analyze mucus thickness and goblet cell numbers in the crypt.

### Immunofluorescence staining and confocal microscopy

For visualization of cell shedding and intracellular bacterial loads, PFA-fixed, OCT-embedded samples of cecum and proximal colon were cryosectioned to 10 μm thickness. Sections were air dried, permeabilized with 0.5% Triton X-100 and blocked with 10% donor goat serum. Sections were stained with anti-Salmonella LPS (Difco; #DF2659-47-5)), anti-Cleaved Caspase-3 (Cell Signaling; #9661S) and anti-Epcam (BioLegend; #118201) antibody. DAPI was used to counterstain nuclei. Tissues were visualized using the Olympus FV3000 microscope.

### Western Blot

For analysis of protein levels specifically in intestinal epithelial cells, colons were harvested and flushed with PBS. The colon epithelial cells were lysed in situ with tissue protein extraction reagent (TPER, Thermo Fisher) containing protease inhibitor cocktail. After 5 min incubation, lysate was recovered, centrifuged at 10,000 g for 3 min to remove debris and stored at −80°C [64]. Whole tissue lysates were obtained by homogenizing tissue samples in 500 μl of TPER containing protease inhibitors and incubating on ice for 20 min. Lysates were centrifuged as above and stored at −80°C. Protein in the samples were quantified using the DC protein assay (BioRad) and approximately 50 μg of protein was loaded for SDS-PAGE. Prestained protein ladder (BioRad) was loaded as a reference. Proteins were transferred on to PVDF membranes and blocked with 4% bovine serum albumin in TBST buffer for one hour. Blots were stained overnight at 4°C using primary antibodies against IL-18 (Abcam; #ab71495) followed by incubation with appropriate secondary HRP-conjugated antibodies. Beta-actin (Santa Cruz; #sc-47778-HRP) levels in the sample were used for normalization. Blots were analyzed using ImageJ software to calculate relative protein levels.

### Dietary intervention

6-8 weeks old stop^flox^ mice were fed regular mouse chow coated with retinyl acetate (500 IU/g) or equivalent amount of vehicle control (corn oil) for 3 days. Chow intake was monitored to ensure both groups of mice consumed similar amounts. Mice were sacrificed and colon lysates were prepared to analyze IL-18 levels using western blot.

### Microscope image acquisition

Image acquisition was performed on the Olympus FV3000 confocal microscope. Images were acquired using a 60X oil immersion lens. For intracellular bacterial data, images were acquired as a Z-stack. All images processing was performed using the Fiji software with an Olympus plugin. Channel color for DAPI was changed to greyscale post-processing. Original 16-bit stacks were converted into RGB format before exporting as a video.

### Statistical analysis

Data shown represent means ± SEM. Data was plotted and analyzed using GraphPad Prism software. For comparison of two groups, Student’s t test was employed. Comparison of two or more groups was performed using One-way ANOVA. Two-way ANOVA was used to compare gene expression across multiple groups.

### Ethics statement

All experiments were approved by and carried out in accordance with the guidelines of the Institutional Animal Care and Use Committee at Brown University (Protocol # 1803000345).

## Acknowledgments

We thank Kevin Carlson for assistance with flow cytometry and Geoff Williams for maintaining the Leduc Bioimaging facility.

## Supporting information

**S1 Fig. Mucus thickness and epithelial turnover analysis.** This figure compares the (A) mucus thickness, (B) goblet cells/crypt and (C and D) epithelial turnover via BrdU incorporation at 2 h (C) and 25 h (D) post injection in homeostatic colons of stop^flox^ and stop^ΔIEC^ mice.

**S2 Fig. Homeostatic colon lamina propria lymphocyte characterization** This figure describes the relative frequencies of (A) CD45+CD3+ cells, (B) CD45+CD3+ IFNγ+ cells, (C) CD45+CD3+IL17+ cells and (D) IL22+ cells in the colons of stop^flox^ and stop^ΔIEC^ mice at homeostasis.

**S3 Fig. Colon lamina propria lymphocyte characterization during infection** This figure describes the relative frequencies of (A) CD3+ IL17+ and (B) CD45+IL22+ cells in colonic lamina propria of stop^flox^ and stop^ΔIEC^ mice 72 hours post *Salmonella* infection.

**S1 Movie. Intracellular *Salmonella* burden in stop^flox^ mice** Video of Z stacks imaging for intracellular loads of *Salmonella* in proximal colon tissues of stop^flox^ mice at 18 hours post infection.

**S2 Movie. Intracellular *Salmonella* burden in stop^ΔIEC^ mice** Video of Z stacks imaging for intracellular loads of *Salmonella* in proximal colon tissues of stop^ΔIEC^ mice at 18 hours post infection.

**S3 Movie. Intracellular *Salmonella* burden in stop^ΔIEC^ + IL-18 mice** Video of Z stacks imaging for intracellular loads of *Salmonella* in proximal colon tissues of stop^ΔIEC^ + IL-18 mice at 18 hours post infection.

