## Supporting information for "Epithelium intrinsic vitamin A signaling co-ordinates pathogen clearance in the gut via IL-18"

**Supporting information file**

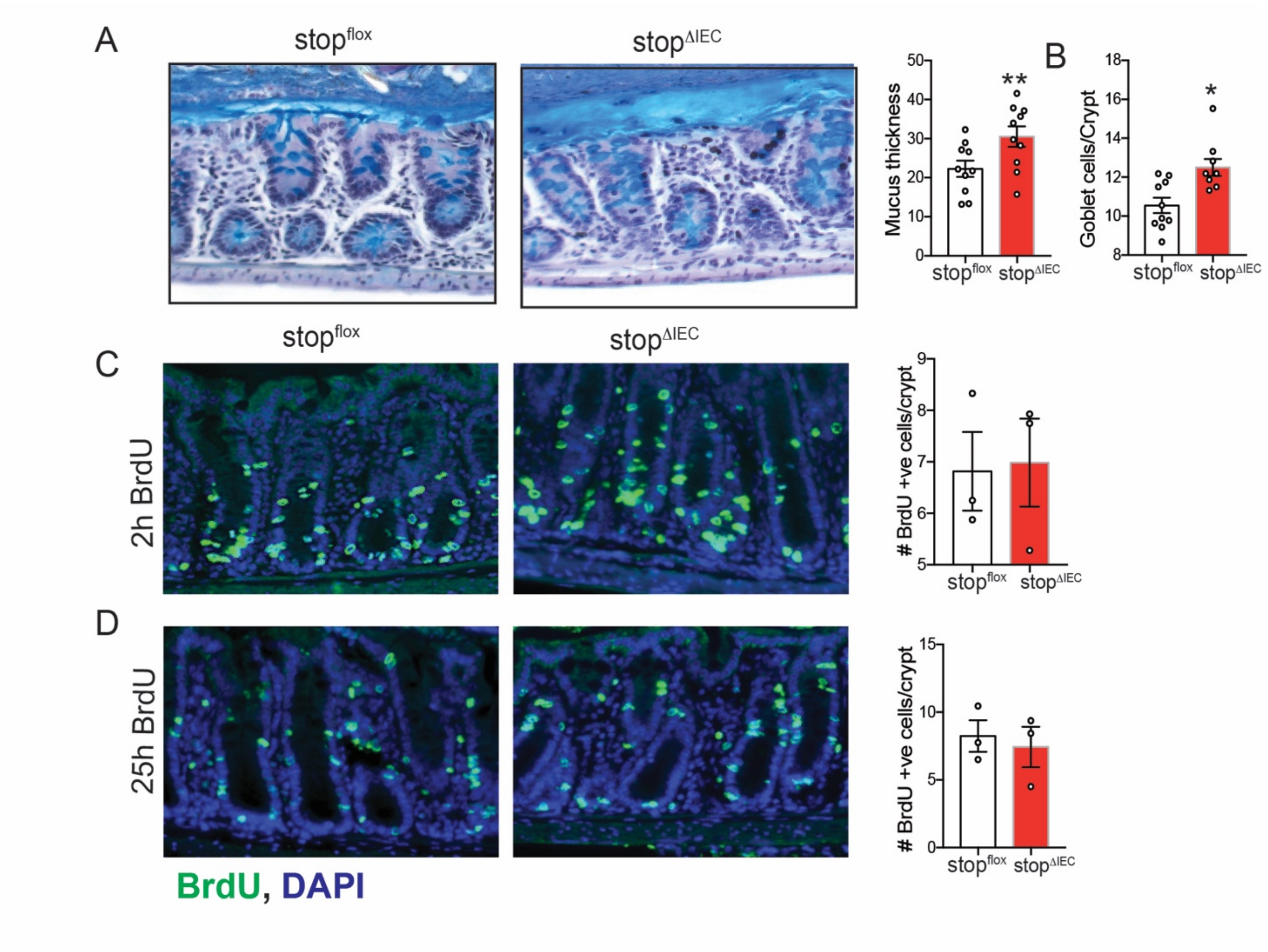


**S1 Fig. Mucus thickness and epithelial turnover analysis**

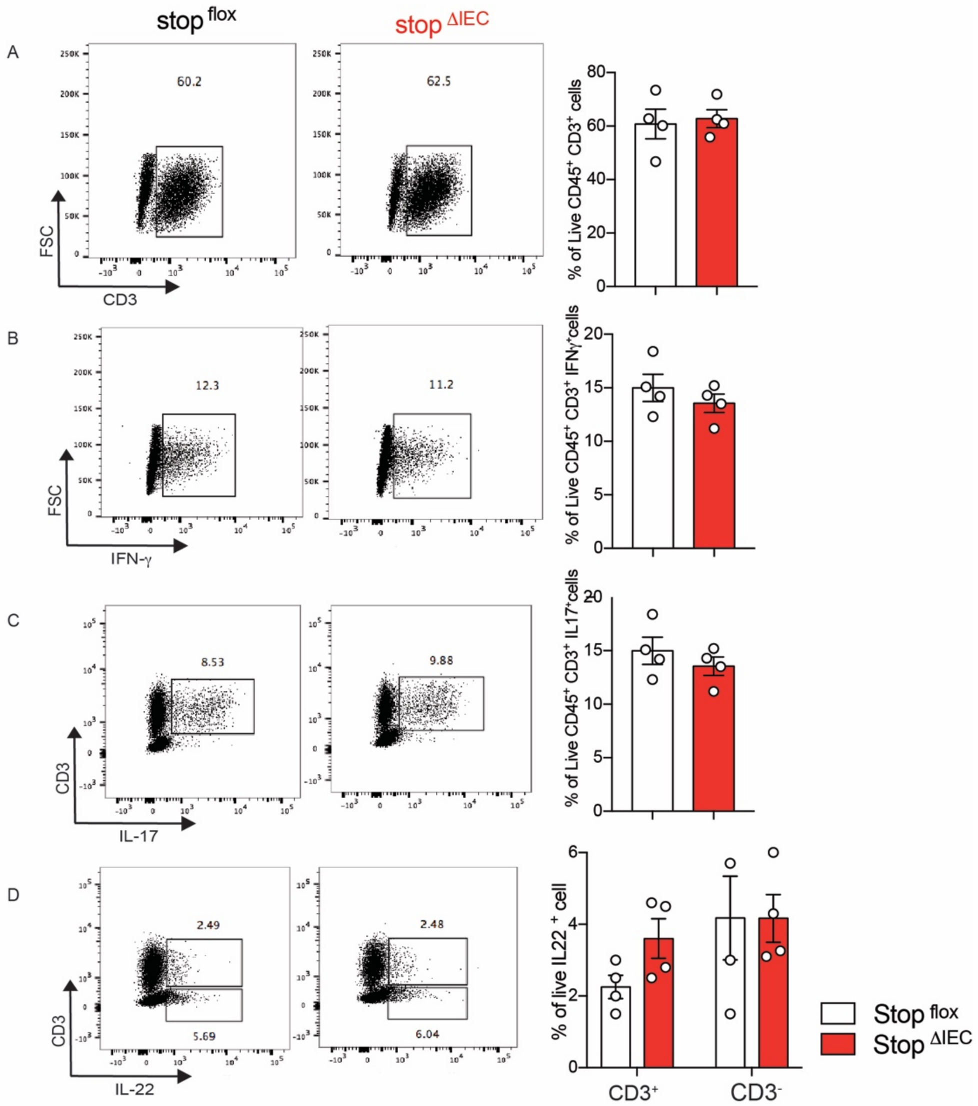


**S2 Fig. Homeostatic colon lamina propria lymphocyte characterization**

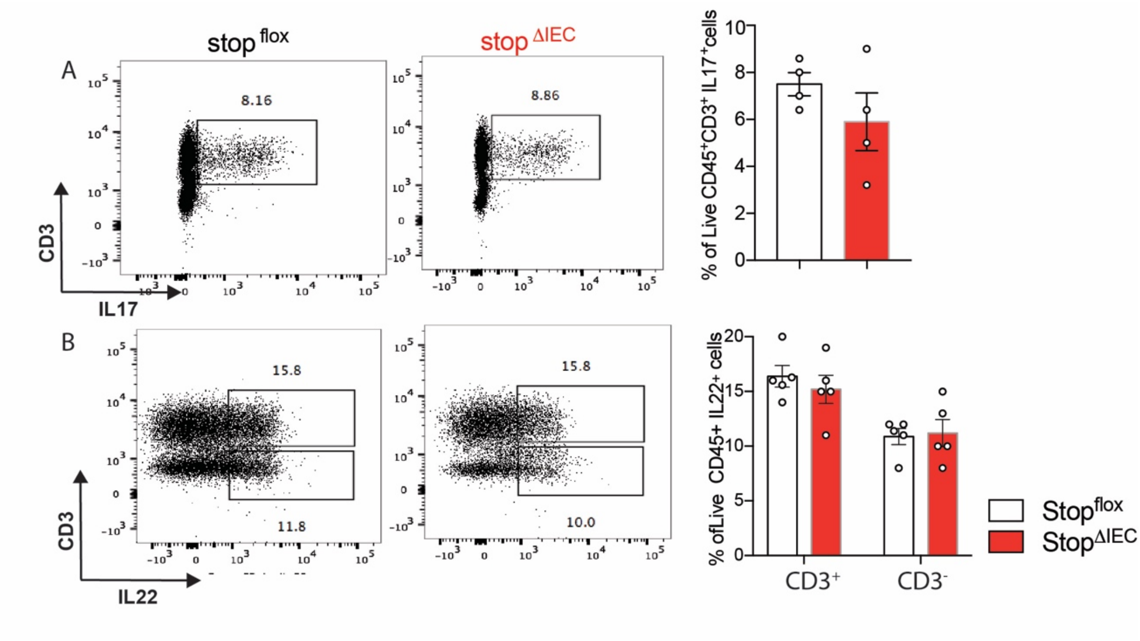


**S3 Fig. Colon lamina propria lymphocyte characterization during infection**
